# Soil respiration in a tropical montane grassland ecosystem is largely heterotroph-driven and increases under simulated warming

**DOI:** 10.1101/501155

**Authors:** Yadugiri V Tiruvaimozhi, Mahesh Sankaran

## Abstract

Soil respiration, a major source of atmospheric carbon (C), can feed into climate warming, which in turn can amplify soil CO_2_ efflux by affecting root, arbuscular mycorrhizal fungal (AMF) and other heterotrophic respiration. Although tropical ecosystems contribute >60% of the global soil CO_2_ efflux, there is currently a dearth of data on tropical soil respiration responses to temperature rise. We set up a simulated warming and soil respiration partitioning experiment in tropical montane grasslands in the Western Ghats in southern India, to (a) evaluate soil respiration responses to warming, (b) assess the relative contributions of autotrophic and heterotrophic components to soil respiration, and (c) assess the roles of soil temperature and soil moisture in influencing soil respiration in this system. Our results show that soil respiration was tightly coupled with soil moisture availability, with CO_2_ efflux levels peaking during the wet season. Soil warming by ~1.4 °C nearly doubled soil respiration from ~0.64 g CO_2_ m^−2^ hr^−1^ on average under ambient conditions to ~1.17 g CO_2_ m^−2^ hr^−1^ under warmed conditions. However, warming effects on soil CO_2_ efflux were contingent on water availability, with greater relative increases in soil respiration observed under conditions of low, compared to high, soil moisture. Heterotrophs contributed to the majority of soil CO_2_ efflux, with respiration remaining unchanged when roots and/or AMF hyphae were excluded. Overall, our results indicate that future warming is likely to substantially increase the largely heterotroph-driven soil C fluxes in this tropical montane grassland ecosystem.

## Introduction

Soils are substantial carbon (C) sinks, storing about 1500 Gt C, ~3 times as in aboveground vegetation (Raich and Schlesinger 1992; Cartmill 2011). They are also significant carbon sources, with soil respiration contributing 64-72 Gt C yr^−1^ to atmospheric pools, which is ~30% of the total terrestrial and marine atmospheric C contribution and ~10 times the C contribution from anthropogenic sources such as fossil fuel combustion (Raich and Schlesinger 1992; Baggs 2006). Tropical ecosystems are estimated to contribute >60% of the global soil CO_2_ efflux (Bond-Lamberty and Thomson 2010; Hashimoto et al. 2015), suggesting that even slight increases in soil respiration levels in these regions can translate to large additions to global atmospheric CO_2_ pools.

Increased atmospheric CO_2_ levels are a major contributor to global warming (IPCC, 2013), which in turn can feed back to influence soil CO_2_ efflux. Many studies have reported warming-induced increases in soil respiration in subtropical, temperate and boreal ecosystems (e.g. Buchmann 2000; Rustad et al. 2001; Conant et al. 2004; Bronson et al. 2007; Lu et al. 2013; Li et al. 2016; Wangdi et al. 2017), and it is estimated that warming has accounted for a 3% increase in soil respiration levels from 1989 to 2008 in tropical ecosystems as well (Bond-Lamberty and Thomson 2010). Warmer conditions can influence CO_2_ fluxes by affecting both autotrophic respiration, from plant roots and plant-associated symbionts such as arbuscular mycorrhizal fungi (AMF), and heterotrophic respiration due to fungal and bacterial decomposers. Temperature rise can lead to altered rates of metabolism in plant roots (Atkin et al. 2000), as well as increased plant C investment in AMF leading to changes in root colonization levels, greater hyphal growth and increased mycorrhizal respiration (Hawkes et al. 2008; Rudgers et al. 2014; Birgander et al. 2017). Heterotrophic respiration can be affected under warmer conditions by changed soil microbial biomass, community composition, bacterial:fungal ratios (Singh et al. 2010; DeAngelis et al. 2015; Auffret et al. 2016), and microbial metabolism leading to altered decomposition rates (see Classen et al. (2015) and references therein for a review of soil microbial (including AMF) responses to warming).

The level of soil respiration contributed by roots, AMF and microbial decomposers, however, can differ across ecosystems. For instance, root respiration can contribute anything between 10 to >90% of the total CO_2_ efflux from soils, while microbial decomposers have been reported to contribute anywhere from 44% up to >70% of the CO_2_ efflux from soils (Atkin et al. 2000; Buchmann 2000; Hanson et al. 2000; Heinemeyer et al. 2012). Ecosystems also differ in soil temperature and soil moisture controls on soil respiration, with CO_2_ efflux variously responding to changes in either or both (e.g. Cartmill 2011; Wu et al. 2011; Liu et al. 2016; Hoover et al. 2016). While there are a number of reports of CO_2_ efflux measurements from tropical ecosystems (e.g. Bond-Lamberty and Thomson (2010) and studies referred therein), there is a paucity of studies that have evaluated soil respiration responses to experimental warming in these ecosystems (Aronson and McNulty 2009; Lu et al. 2013). Consequently, there is a dearth of data on tropical soil respiration responses, relative contributions of the autotrophic and heterotrophic components, and abiotic controls on CO_2_ efflux under temperature rise.

We evaluated soil respiration responses to simulated warming in a tropical montane grassland ecosystem in the Western Ghats biodiversity hotspot, India, and assessed the relative contributions of roots, AMF and decomposers to soil CO_2_ efflux, over ~2 years. These montane grasslands support high biodiversity but are also threatened as land use change has greatly reduced their extent, and the remaining grasslands are believed to be particularly vulnerable to climate change (Sukumar et al. 1995; Arasumani et al. 2018). In particular, we tested the prediction that soil respiration will be higher under simulated warming than under ambient (control) conditions. In addition, we quantified autotrophic and heterotrophic contributions to soil respiration in this ecosystem by measuring CO_2_ efflux with and without plant roots and/or AMF hyphal components. We also assessed the roles of soil temperature and soil moisture in influencing instantaneous soil respiration in this ecosystem.

## Methods

### Study area

The experiment was conducted in tropical montane grasslands of the shola-grassland ecosystem, a unique mosaic of grassland interspersed with pockets of stunted evergreen forests (sholas), found in the higher reaches of the Western Ghats (Robin and Nandini 2012). These grasslands are representative of other montane grassland and forest-grassland mosaics globally, such as Afromontane ecosystems (Kotze and Samways, 2001; Parr et al. 2014), Campos-Araucaria forest mosaics in southern Brazil (Overbeck et al. 2007), and forest-patana grassland mosaics in Sri Lanka (Gunatilleke et al. 2008).

Our experiment was located in the Avalanche area of the Nilgiris Biosphere Reserve (11.27° N 76.55° E, elevation: ~2300 m), in the state of Tamil Nadu in southern India. The average annual temperature in the region is 14.4 °C, and the average annual rainfall is 1847 mm (https://en.climate-data.org/location/24046/). The majority of the precipitation in these grasslands occurs during the South-West monsoon season from early June to early September, and the North-East monsoon season from early October to early December, accounting for ~905 mm and ~528 mm rainfall on average, respectively. Summer precipitation from early March to late May averages ~200 mm, while the winter months from late December to late February are the driest (District Statistical Handbook, The Nilgiris, 2015-2016; http://nilgiris.nic.in/images/districthandbook0809.pdf). Temperatures peak around May (average temperature: 16.6 °C), and are lowest around January (average temperature: 12.4 °C) (https://en.climate-data.org/location/24046/), with winter temperatures frequently going below 0 °C.

These grasslands support several species of grasses and herbs, and a variety of wild herbivores such as sambar (*Rusa unicolor*), gaur (*Bos gaurus*), Asiatic elephant (*Elephas maximus*) and the endemic Nilgiri tahr (*Nilgiritragus hylocrius*).

### Experimental setup

#### Open top chambers (OTCs)

To study soil respiration responses to simulated temperature rise, we used 9 open top chambers (OTCs), which are passive warming structures (Aronson and McNulty 2009; Godfree et al. 2011), and adjacent paired control plots experiencing ambient temperature conditions. Three OTCs and control plots each were set up within three 10 m × 20 m fences located in areas of similar slope and aspect, in May and June 2014. The OTCs were hexagonal structures, ~3 m in diameter and ~50 cm tall (design modified from Godfree et al. (2011)). Iron frames supported the pyramidal structures, each with five sides having acrylic/polycarbonate walls placed at an inclination of ~60° to the ground and the sixth side left open after initial trials at our field site showed us that OTCs with all six sides closed increased temperatures up to ~11 °C, as opposed to up to ~4 °C in the 5-sided OTCs (data not shown). The control plots were 1 m × 1 m in dimension.

#### Respiration partitioning treatments

To assess autotrophic and heterotrophic contributions to soil respiration in our system and to measure their responses to warming, we set up respiration collars within the OTCs and adjacent to the control plots. Soils within these collars were ‘partitioned’ to measure respiration contributions of ‘full soil’, soil without roots, and soil without roots and AMF (referred to here on as ‘partitioning treatments’; protocol adapted from Marthews et al. 2014; Fig. 1). Each OTC/control plot had three collars, one each for the three soil partitioning treatments, for a total of 54 collars. The partitioning treatments were set up in the first week of November 2014.

**Fig. 1.**
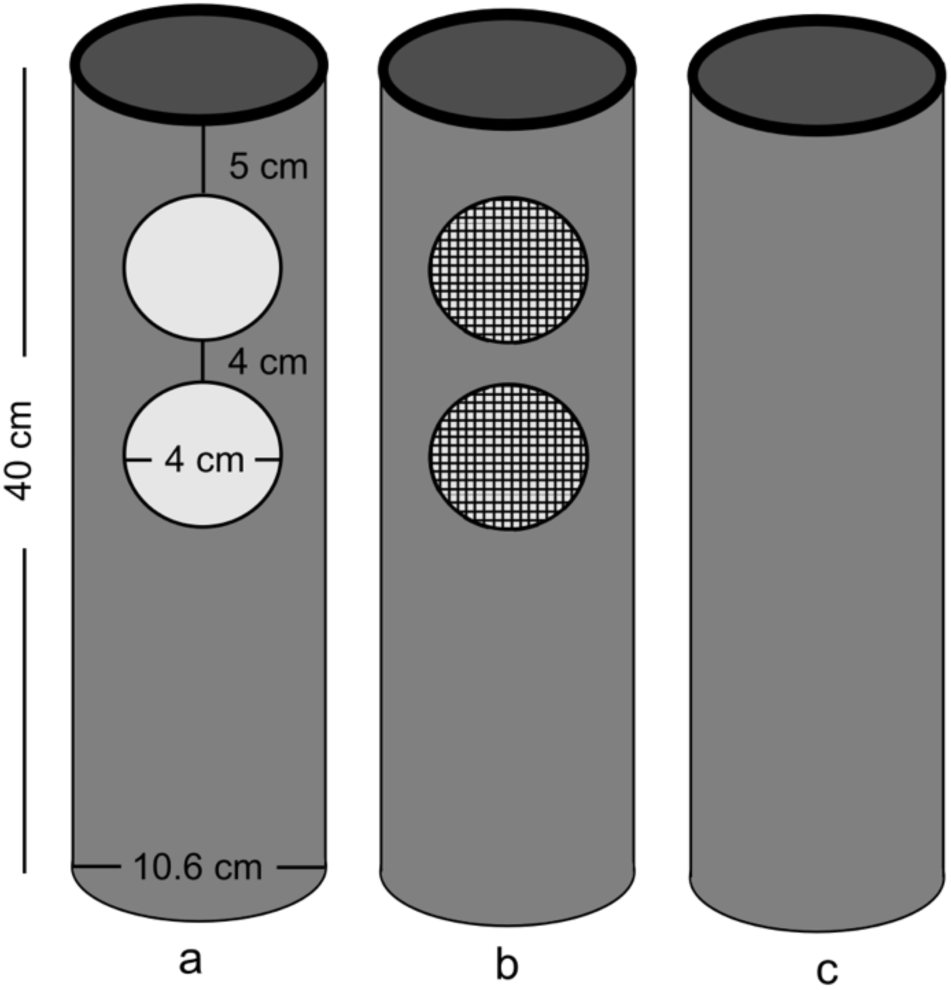
A representation of the design and dimensions of the PVC pipe collars used for the soil partitioning treatments. (a) Full soil treatment, with circles representing the holes made on one side of the collars, (b) Soil without roots treatment, with circles representing holes covered by nylon meshes to keep out fine roots, and (c) Soil without roots and AMF treatment.

Briefly, the partitioning treatments consisted of polyvinyl carbonate (PVC) pipe collars (length: 40 cm, diameter: 10.6 cm) buried in the soil to a depth of ~35 cm with ~5 cm remaining above the ground. Pits ~35 cm deep were made at each collar location, and were re-filled with the same soil after inserting the collar and sifting through the soil to remove all severed roots and other organic debris. The ‘full soil’ treatment had collars with 4 circular holes of ~4 cm diameter, with 2 pairs of holes drilled along opposite sides of the collars (Fig. 1a). These holes allowed roots and AMF hyphae in the top ~12 cm of the soil to freely grow into the collars, and contribute to CO_2_ efflux. Collars for the ‘soil without roots’ treatment had similar holes drilled, but covered with nylon meshes with 40µ pores to allow the growth of AMF extraradical mycelium (ERM) into the collars, but not fine roots, so that CO_2_ efflux measured from these collars do not have the root contribution (Fig. 1b). The ‘soil without roots and AMF’ treatment had collars with no holes, preventing the growth of roots or AMF hyphae into them, thus allowing for CO_2_ efflux measurement from root- and AMF-free soil (Fig. 1c).

We tested the efficacy of the partitioning treatments using additional treatment collars set up in early October 2015, from which we collected soils and measured the amounts of roots, AMF hyphae and microbial biomass in them in November 2016. We found that the treatments were successful in allowing/preventing the growth of roots and/or AMF hyphae within the collars (detailed methods and results in Supplementary Information A). Further, to ensure that soil disturbance during the setting up of the collars did not affect soil respiration over the period of our study, we set up two types of ‘method control’ collars that were installed along with the treatment collars, within an OTC-control plot pair in each fence. One of these (designated as C1) consisted of PVC collars set up exactly as the other treatment collars, but without removing severed roots and other debris from the soils before filling in the collars after installation. The second (designated as C2) had PVC collars hammered into the ground to a depth of ~30 cm without displacing the soil before installation. We found that, after an initial spike, soil respiration in these ‘method control’ collars was indistinguishable from the ‘soil without roots and mycorrhizae’ treatment (details of methods and results in Supplementary Information A). We therefore report only results from the three partitioning treatment collars.

### Soil respiration measurement and calculations

Soil respiration in the partitioning treatment and control collars were estimated at ~15 day intervals from late November 2014 to late January 2017, for a total of 48 sampling days. We used a portable IR-based gas analyser (Environmental Gas Monitor; EGM-4, PP Systems, USA), to measure CO_2_ flux. Alongside CO_2_ flux measurements, we also measured ambient atmospheric temperature within each OTC and control plot, and collar height (averaged across three measurements per collar) at each measurement time point. These data were then used to calculate CO_2_ efflux following Marthews et al. (2014), as:

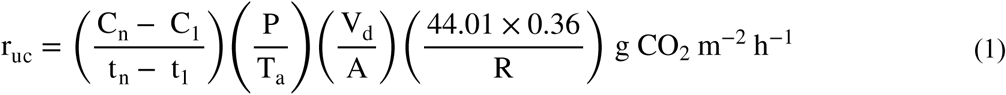

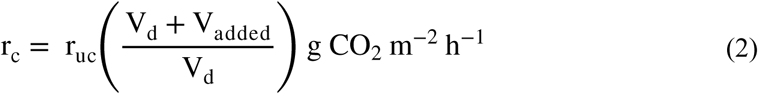

where r_uc_ denotes soil CO_2_ efflux calculated without correcting for the added volume of the respiration collar, and r_c_ denotes CO_2_ efflux corrected for volume; C_n_ - C_1_ is the CO_2_ flux difference, typically between the last 10 readings per measurement, or between the first and last flux values if the measurement had less than 10 readings; t_n_ - t_1_ is the difference in time, in seconds, over which the difference in CO_2_ flux was calculated; is ambient atmospheric pressure, in mb, averaged over t_n_ - t_1_ as measured by the EGM; T_a_ is atmospheric temperature in Kelvin; V_d_ is volume within the EGM respiration chamber; is the area of soil over which CO_2_ flux was measured; R is the Universal Gas Constant, 8.314 J K^−1^ mol^−1^; and V_added_ is the volume of the respiration collar above the soil surface at the time of measurement. A more detailed discussion of the method for measurement and calculation of CO_2_ efflux can be found in Marthews et al. (2014).

To account for potential measurement errors in CO_2_ flux while using the EGM in the field, we assessed linearity of CO_2_ accumulation for each collar for each sampling day using linear models of CO_2_ accumulation versus time, and only those measurements that satisfied the criteria that R^2^ ≥ 0.9 (Savage et al. 2008), and had a positive slope, were used for further analyses. Further, since some values of measured CO_2_ efflux were found to be unusually high or low, we excluded values that fell beyond 3 SDs of the mean CO_2_ efflux from the analyses. Our final analyses were based on 80.9% of the original data collected.

### Temperature and soil moisture measurements

We used iButtons (Thermochron Temperature Data Loggers, Maxim Integrated, USA) to measure air and soil temperatures. Air temperatures were measured from December 2014 to January 2017 by placing iButtons ~2-3 cm above the ground within all 9 OTCs, and 3 control plots, one in each fence. In addition, we also measured soil temperatures from May 2015 to January 2016, by placing iButtons just below the soil surface in an OTC and control plot in each fence. Data for some months are missing due to loss of iButton, or due to logging errors. We also measured instantaneous soil temperature and soil moisture when quantifying soil respiration. Instantaneous soil temperature measurements were done at 12.5 cm depth using a temperature probe (HI145-00 and HI145-01, Hanna Instruments, USA), with ~3 replicates in the vicinity of each soil respiration collar. Soil moisture measurements were done over the top 12 cm of soil using a soil moisture meter (FieldScout TDR 100 Soil Moisture Meter, Spectrum Technologies, USA), with ~3 replicates in the vicinity of each soil respiration collar.

### Data analysis

We used linear mixed effects models (LMMs) to test whether (a) soil respiration in warmed plots differed from control plots, (b) CO_2_ efflux was different in treatments where roots and/or AMF were excluded from ‘full soil’ respiration levels, and (c) partitioning treatment effects were different in warmed versus control plots. Warming (OTC/control), partitioning treatments (full soil, soil without roots, soil without roots and AMF) and their interaction were used as fixed factors and collars nested within plots, which were in turn nested within fences were used as random factors to account for multiple measurements across time from the same respiration collars (Baayen et al. 2008; Zuur 2009; Cunnings and Finlayson 2015). Soil respiration, measured at ~15 day intervals from November 2014 to January 2017 per warming and partitioning treatment, was the response variable. In all, there were 310-348 individual respiration measures per partitioning treatment per OTC/control plot (median = 339.5), for a total of 1994 values. Soil CO_2_ efflux values were log transformed before analyses to meet model assumptions.

Further, we assessed whether instantaneous soil temperature and soil moisture individually or together best explained variation in soil respiration in the OTC and control plots using LMMs in conjunction with AIC based model selection. Instantaneous soil temperature and soil moisture, warming treatment and their interactions were as fixed factors and as in the previous analysis, collars nested within plots, which were in turn nested within fences were the random factors. We only used soil respiration and instantaneous soil moisture and soil temperature data from the ‘full soil’ collars for these analyses. The fixed effects in the three candidate LMMs were (a) soil temperature × soil moisture × warming treatment, (b) soil temperature × warming treatment, and (c) soil moisture × warming treatment. We also computed marginal and conditional R^2^ values that give an indication of the variation explained by only the fixed effects and the fixed and random effects, respectively, for all three candidate models (Nakagawa and Schielzeth 2013). Again, soil CO_2_ efflux values were log transformed before analyses to meet model assumptions.

We used the R package *lme4* to conduct all the mixed effects models (Bates 2010; Bates et al. 2014, Kuznetsova et al. 2015; Bates et al. 2017), the *lmerTest* package to conduct *t*-tests using Satterthwaite approximations for the degrees of freedom, and the *car* package to conduct Type II Wald chi-square tests to assess the statistical significance of the fixed effects (Bates et al. 2014, Bates et al. 2017). Marginal and conditional R^2^ values were computed using the *piecewiseSEM* package (Lefcheck 2016). All analyses were conducted using R version 3.2.4 (The R Foundation for Statistical Computing, 2016).

## Results

### Soil and air temperature and soil moisture status in the OTC and control plots

Warming treatment effects on soil temperature were marked, with average daily soil temperatures of control plots and OTCs at 16.10 ± 0.13 °C and 17.53 ± 0.15 °C (over 277 measures from 146 days each), respectively. The difference in mean daily temperature between OTC and control plots averaged 1.41 ± 0.08 °C overall (Fig. 2a), with a maximum temperature increase of 3.43 °C. Monthly averages of soil temperatures ranged from 14.55-21.67 °C in the controls and 16.09-22.93 °C in the OTCs (Fig. 2b).

**Fig. 2.**
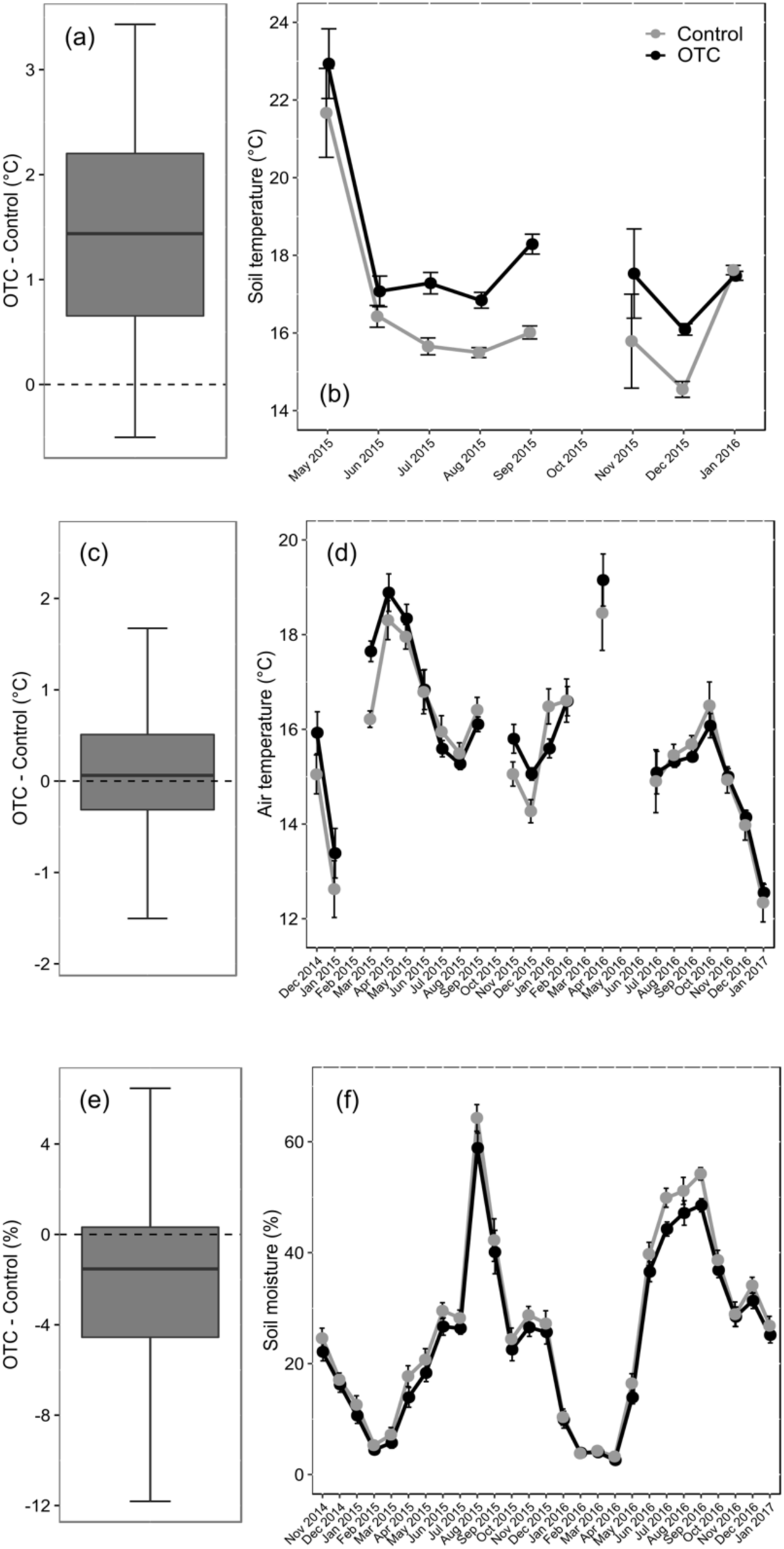
Overall daily average differences between OTCs and control plots for (a) Air temperature (c) soil temperature and (e) soil moisture, and (b), (d) and (f), the above three variables averaged per month, respectively, in the control and OTC plots. Dots in grey denote control plots and those in black denote OTCs. Error bars in (b), (d) and (f) are 1SE around the mean.

Over the period of the study, average daily air temperatures of control plots and OTCs were 15.65 ± 0.08 °C (from 793 measures over 371 days) and 15.42 ± 0.05 °C (from 1985 measures over 371 days), respectively. However, the difference in mean air temperatures of OTCs and control plots per day averaged 0.10 ± 0.04 °C (Fig. 2c), with a maximum air temperature increase in the OTCs of 2.62 °C. Average monthly air temperatures ranged 12.34-18.46 °C in the controls and 12.55-19.15 °C in the OTCs (Fig 2d).

Soil moisture was lower in the OTCs than in the control plots (Fig. 2e), with averages of 23.74 ± 0.76% and 25.98 ± 0.81%, respectively (from 421 and 429 estimates, respectively, across 27 months). Soil moisture levels ranged from 3.26-64.30% and 2.60-58.88% on average in the controls and OTCs, respectively (Fig. 2f).

### Effects of warming and partitioning treatments on soil respiration

Warming significantly increased soil respiration by ~55-89% compared to control levels in all three partitioning treatments (*P* < 0.05). The three partitioning treatments, however, were statistically indistinguishable from each other in both controls and OTCs (Table 1, Fig. 3). Average soil respiration over the entire duration of the experiment across partitioning treatments was 0.62±0.01 g CO_2_ m^−2^ hr^−1^ in the controls and 1.16±0.03 g CO_2_ m^−2^ hr^−1^ under warmed conditions.

**Table 1.** Summary of LMM results of soil respiration responses to warming treatment, partitioning treatments, and instantaneous soil temperature and soil moisture.

| Effect | Wald chi-square | df | *P* |
| --- | --- | --- | --- |
| Warming | 44.7641 | 1 | <0.001 |
| Partitioning treatment | 2.5216 | 2 | 0.28 |
| Warming × Partitioning treatment | 1.1280 | 2 | 0.57 |
| Soil moisture* | 57.8397 | 1 | <0.001 |
| Soil temperature | 0.0005 | 1 | 0.98 |
| Warming | 9.7556 | 1 | 0.002 |
| Soil temperature × Soil moisture | 12.7965 | 1 | <0.001 |
| Soil temperature × Warming | 0.1610 | 1 | 0.69 |
| Soil moisture × Warming | 2.8782 | 1 | 0.09 |
| Soil temperature × Soil moisture × Warming | 2.0987 | 1 | 0.15 |
| Soil moisture† | 57.1421 | 1 | <0.001 |
| Warming | 9.1714 | 1 | 0.002 |
| Soil moisture × Warming | 5.1778 | 1 | 0.023 |

\* Values in this section are from the ‘full model’, an LMM with soil moisture, soil temperature, warming treatment and interactions as fixed effects.
† Values in this section are from the ‘best model’ (see Table 2), an LMM with soil moisture, warming treatment and their interactions as fixed effects.

**Fig. 3.**
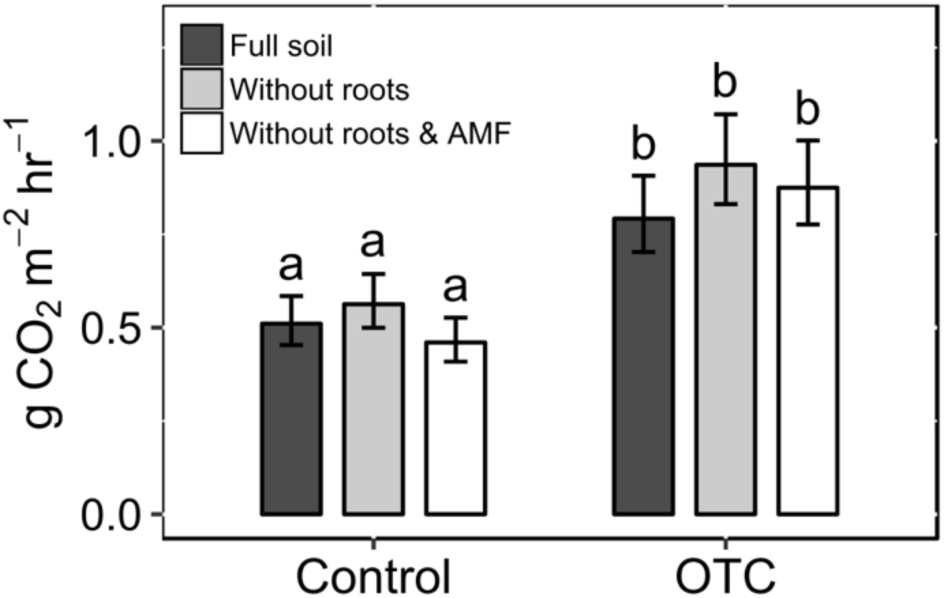
Average overall soil respiration in control and warmed conditions in the three partitioning treatments. ‘Full soil’ treatment is represented by dark grey bars, ‘soil without roots’ by light grey bars, and ‘soil without roots and AMF’ by white bars. Error bars represent 1SE around the mean, obtained from the mixed effects model used for analysis. Different letters indicate statistically significant differences among treatments (*P* < 0.01 or lesser).

Soil respiration also differed by month mirroring the seasonality of our study system (Fig. 4). Overall, control plots recorded minimum soil respiration of 0.16±0.02 g CO_2_ m^−2^ hr^−1^ in March 2015, while respiration peaked to 1.10±0.12 g CO_2_ m^−2^ hr^−1^ in September 2016; while the OTCs recorded minimum and maximum soil respiration levels of 0.63±0.07 g CO_2_ m^−2^ hr^−1^ and 1.84±0.36 g CO_2_ m^−2^ hr^−1^, in February 2015 and April 2016, respectively (Fig. 4). Responses per partitioning treatment were similar to this overall warming effect (Supplementary Information B).

**Fig. 4.**
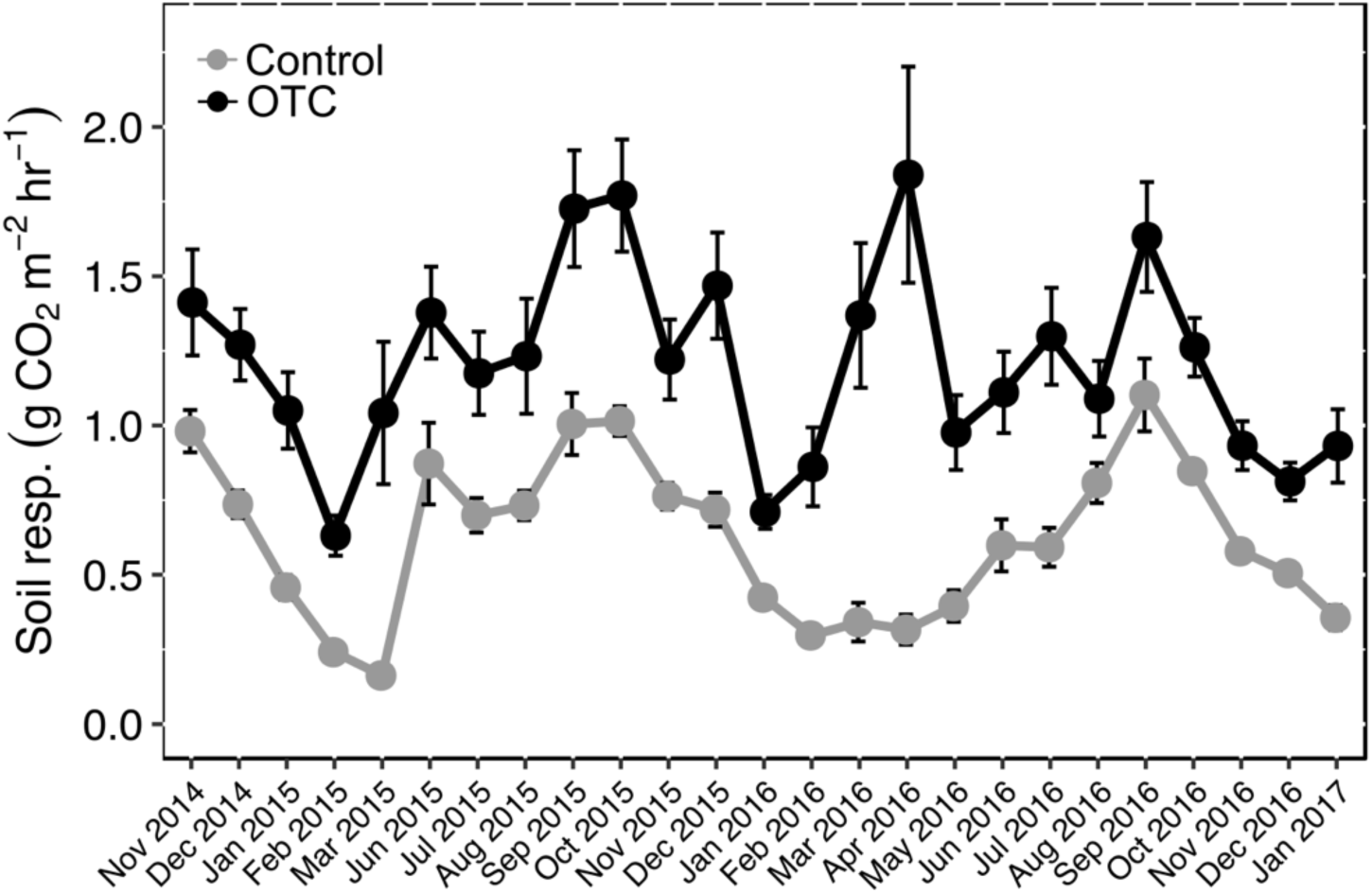
Average soil respiration in control and OTC plots averaged per month, across partitioning treatments. Dots in grey represent control plots and those in black represent OTCs. Error bars represent 1SE around the mean.

### Effects of soil temperature and moisture on soil respiration

The most parsimonious model predicting soil respiration in our system was the model that included warming treatment, instantaneous soil moisture and their interaction as predictors (Table 2). Soil respiration was higher in the warmed plots (*P* = 0.002; Table 1), and increased as soil moisture increased (*P* < 0.001; Table 1). However, the effect of warming on soil respiration was more pronounced under low soil moisture conditions (soil moisture × warming: *P* = 0.023; Table 1; Fig. 5). Given that the partitioning treatments were statistically indistinguishable, only data from the ‘full soil’ treatment were used for these analyses.

**Table 2.** Model comparisons of LMMs to assess instantaneous soil moisture and/or temperature and warming treatment effects on soil respiration. Marginal and conditional R^2^ values give an indication of the variation explained by the fixed effects only and the fixed and random effects together, respectively, in mixed effects models.

| Model (fixed effects) | AIC | $\Delta AIC$ | $R^2_{\text{marginal}}$ | $R^2_{\text{conditional}}$ |
| --- | --- | --- | --- | --- |
| Soil moisture × Soil temperature × Warming | 1455.884 | 28.524 | 0.16 | 0.30 |
| Soil temperature × Warming | 1478.419 | 51.059 | 0.06 | 0.22 |
| Soil moisture × Warming | 1427.360 | 0 | 0.15 | 0.28 |

**Fig. 5.**
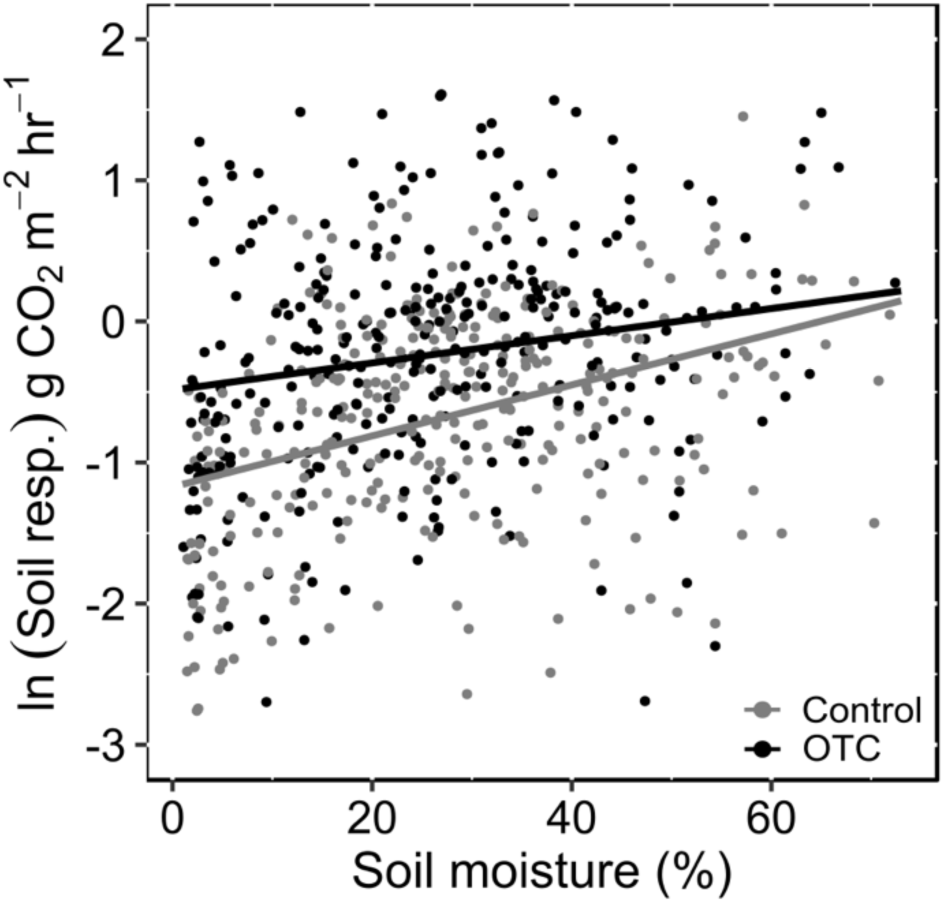
Soil respiration under control and warmed conditions in the ‘full soil’ treatment are positively related to average soil moisture. Control: ln(CO_2_ efflux) = −1.17 + (0.02 × soil moisture); OTC: ln(CO_2_ efflux) = −0.49 + (0.01 × soil moisture). Both slopes are significantly different from 0 (*P* < 0.001 in both cases). Slopes were obtained from linear mixed effects model analysis. Data for control plots are represented in grey and OTCs in black.

## Discussion

Results from this study suggest that warming drastically increases soil respiration in this tropical montane grassland ecosystem in the Western Ghats. Soil respiration was largely heterotrophic, with exclusion of roots and/or AMF hyphae not significantly changing CO_2_ efflux levels. Further, soil respiration increased with soil moisture in these grasslands, with CO_2_ efflux maxima and minima coinciding with the end of the wet and dry seasons, respectively. However, warming effects on soil respiration were more pronounced in drier soils.

Soil respiration in the study site was largely heterotroph-driven. Heterotroph-dominant soil respiration, as observed in this study system, has been reported from several non-forest ecosystems, such as grasslands, croplands and oak savannas, and also some temperate forests, although autotrophs generally contribute ~50% of the soil respiration in forest ecosystems (Kelting et al. 1998; Buchmann 2000; Hanson et al. 2000; Melillo et al. 2002; Scott-Denton et al. 2006; Cartmill 2011). Heterotrophic contributions to soil respiration have been shown to correlate strongly with soil detritus levels (Bond-Lamberty et al. 2004) and increase with increasing soil nitrogen availability (Rodeghiero and Cescatti 2006). Heterotroph contributions to soil respiration are also potentially influenced by other factors such as vegetation and soil microbial community composition, net primary production and litter quality, though which of these factors underlie heterotroph dominated respiration in these montane grasslands is unclear.

Another potential reason for the lack of differences in soil respiration levels among partitioning treatments in this study is soil microbial biomass changes following loss of plant roots and AMF hyphae. Plants and microbes compete for soil resources (such as N), and removal of plant roots can lead to increases in decomposer biomass, which in turn can increase soil CO_2_ efflux levels, masking the loss of the autotrophic contribution to soil respiration. In this study, while microbial biomass carbon (MBC) increases with root exclusion, there are no increases in microbial biomass with AMF hyphal exclusion (Supplementary Information A, Fig. A.1). This suggests that the lack of differences in soil respiration levels between treatments in this study is not entirely a consequence of decomposer biomass increases in the root- and AMF-free soil. In other words, MBC responses to root and AMF hyphal exclusion in this study further supports the conclusion that soil respiration in our system is heterotroph dominated.

Soil CO_2_ efflux levels in this experiment nearly doubled, from ~0.62 g m^−2^ hr^−1^ under ambient conditions to ~1.16 g m^−2^ hr^−1^ within the OTCs. This is in agreement with empirical findings from other ecosystems including grasslands, as well as theoretical studies suggesting increases in soil respiration under climate change in the tropics (Schindlbacher et al. 2009; Bond-Lamberty and Thomson 2010; Lu et al. 2013; Wang et al. 2014). The observed increases in respiration under warmer conditions can be driven by several mechanisms. First, greater soil CO_2_ efflux can result from greater microbial metabolism under warmed conditions, given that heterotrophs contribute the majority of the respiration in this system (Schindlbacher et al. 2011; Luo et al. 2014). Previous studies have also shown that autotroph and heterotroph respiration responses to changing temperature regimes can be very different, with heterotrophs, rather than autotrophs, reported to be more sensitive to increasing temperatures (Wei et al. 2010; Li et al. 2013; Wang et al. 2014). Second, higher levels of labile C under warmed conditions can drive shifts to a more ‘rapid’ nutrient and C cycling system, leading to greater soil respiration (Metcalfe et al. 2011; Luo et al. 2014). Soil respiration responses to warming can also be mediated by microbial community shifts, and soils with different microbial community compositions have been demonstrated to respond differently to temperature increases (Auffret et al. 2016). For instance, warming has been shown to promote certain bacterial phyla over fungal phyla (Luo et al. 2014), and increased bacteria:fungi ratios are associated with faster C and nutrient cycling (Wardle et al. 2004). Fungal communities, too, have been demonstrated to shift under warming to favour taxa that are better decomposers of recalcitrant C (Treseder et al. 2016), which would then amplify CO_2_ efflux from these soils. However, at present, it is not clear which of these mechanisms may be driving warming-mediated increases in soil respiration in our study system.

Soil respiration was positively related to soil moisture in these markedly seasonal montane grasslands, with clear wet and dry seasons. Moisture effects on (especially heterotrophic) soil respiration have been widely reported, and function by affecting several physiological, biochemical and ecological factors such as decomposer substrate availability, nutrient and dissolved organic matter mobility, osmoregulation and changes in microbial community composition (Orchard and Cook 1983; Scott-Denton et al. 2006; Wei et al. 2010; Yan et al. 2009; Moyano et al. 2013). Further, experiments in temperate ecosystems have demonstrated that while soil respiration minima correspond to low temperature conditions, peaks coincide with the ‘growing season’, often responding to moisture rather than temperature maxima (Heinemeyer et al. 2012; Hoover et al. 2016; Liu et al. 2016). While soil respiration peaked in the wet season in this study system, warming amplified soil respiration during the dry season. Warming-mediated amplification of respiration during the drier months can be because some of the driest months in these montane grasslands are also the coldest, during which soil microbes under warmed conditions will likely have greater metabolism leading to the higher levels of respiration. Overall, we see the highest respiration levels under wetter and warmer conditions. Indeed, a modelling study analysing global soil respiration responses to environmental factors also suggests that the regions with high soil respiration, globally, are associated with both high temperature and precipitation (Hashimoto et al. 2015).

On the whole, the present study indicates that warming is likely to substantially increase soil respiration levels in this tropical montane grassland ecosystem, with effects more pronounced under drier conditions. While the mechanisms behind soil respiration responses to warming in our study system are as yet unclear, our results suggest that decomposers play a major role in regulating observed soil CO_2_ efflux responses to warming. In the longer term, acclimation over time of roots, AMF and other soil components to altered temperature regimes, or depletion of resources such as water or labile carbon, might alter CO_2_ efflux responses to warmer temperatures (Atkin et al. 2000; Luo et al. 2001; Melillo et al. 2002; Kirschbaum 2004; Heinemeyer et al. 2006; Auffret et al. 2016; Romero-Olivares et al. 2017). Soil respiration can also be affected in the long term by warming-mediated alteration of factors such as vegetation composition and structure (Cartmill 2011; Metcalfe et al. 2011; Rudgers et al. 2014; Mayer et al. 2017), length of growing season (Rustad et al. 2001), AMF species pool (Kim et al. 2015) and decomposer community composition (Zogg et al. 1997; DeAngelis et al. 2015). Other global change factors, such as increased atmospheric nutrient deposition, can also influence warming effects on plants and microbes (Olsson et al. 2005). Future longer term studies that also estimate warming-induced changes in other parameters such as vegetation growth, foliar respiration, soil microbial biomass and other components of soil C are needed to assess the net contribution of these ecosystems as sources or sinks of carbon.

## Acknowledgements

We are very grateful to Suseelan K, A Selvakumar, Siva and Manikantan, whose help made the experiment possible. We also gratefully acknowledge several past and current members of the BEER lab at NCBS, especially Siddharth Iyengar, Chandan Pandey, Atul Joshi, Dincy Mariyam, Harinandanan P V, and Chengappa S K, and many residents of Emerald village in the Nilgiris for their generous help with field work. We thank the Tamil Nadu Forest Department for permission to conduct the experiment. The Rufford Foundation and National Centre for Biological Sciences provided funding for this study.

## Supplementary information A - Testing the efficacy of the partitioning treatments and evaluating the effect of soil disturbance during experimental setup on soil respiration

### Methods

To measure the efficacy of the soil partitioning treatments in allowing/preventing the growth of roots and/or arbuscular mycorrhizal fungal (AMF) hyphae into the collars, 9 additional collars, one for each partitioning treatment in each of the three fences, were set up in early October 2015, outside the open top chambers (OTCs) and control plots. We harvested soil from these collars in November 2016 and transported the samples to the National Centre for Biological Sciences, Bangalore, to measure root biomass, AMF extraradical mycelial (ERM) length and soil microbial biomass. The samples were stored at ~4 °C till they were analysed.

To measure root biomass, soils from these samples were sifted to extract roots, which were then dried at 60 °C for 48 hr and weighed.

AMF ERM lengths in all samples were measured following Brundrett and others (1994). Briefly, ~5g of each airdried soil sample was suspended for 30 min in dilute sodium hexametaphosphate (Calgon) solution, aliquots of which were then passed through a 20µ nylon membrane on a vacuum filter to extract hyphae. These were then re-suspended in and incubated for 1.5 hr in trypan blue stain. Stained hyphae were extracted onto gridded 1.2µ cellulose nitrate filters, which were then air dried, placed on glass slides, cleared with low viscosity, low fluorescence immersion oil and observed under 400× magnification. Intersections of stained aseptate hyphae with a 10 × 10 grid on an eyepiece graticule were counted over 50-65 (median = 55) microscopic fields of view scanning across each sample slide, and hyphal length per slide was calculated. A subsample (~3 g) of each soil sample was used to measure gravimetric water content (as the difference in soil subsample weight before and after drying at 110 °C for 48 hr, per g dry weight of soil), and these data, along with the volume of solution that was used to dilute the samples, were used to calculate AMF ERM length per g dry weight of soil (Brundrett and others 1994).

Microbial biomass carbon (MBC) was estimated using substrate induced respiration and titration (method adapted from Anderson & Domsch 1978; Höper 2006; Fanin and others 2011). Briefly, 10 g dry weight equivalent of each air dried soil sample was pre-incubated at 30 °C at near 80% water holding capacity (WHC) for 72 hr in airtight plastic containers. Glucose (1.6 g per g dry soil) was then added as solution to the samples, bringing the soils up to 80% WHC, immediately after which a vial with 2 ml 2N NaOH was placed in the containers to serve as the alkali CO_2_ trap. The sealed containers were then incubated for 24 hr at 30 °C. Two airtight containers without soil samples but with vials containing the NaOH trap were also kept for incubation to serve as controls. After incubation, NaOH in the traps were titrated with phenolphthalein indicator against 0.5N HCl to estimate CO_2_ release, which was then used to calculate MBC per kg dry weight of soil (Höper 2006).

In order to ascertain that soil displacement and handling during partitioning treatment setup did not affect soil respiration over the period of our study, two types of ‘method control’ collars were installed along with the treatment collars. One of these (designated as C1) consisted of 40 cm PVC pipes without holes, inserted to ~35 cm depth, similar to the treatment collars, but without sifting through the soil to remove roots and organic debris. The second (designated as C2) consisted of similar PVC pipes hammered into the soil (to a depth of ~30 cm), without displacing the soil prior to installation. Respiration measurements from these controls were expected to be similar to the ‘soil without roots and AMF’ treatment, after an initial spike in CO_2_ efflux as severed roots and other debris are decomposed. A collar each for the C1 and C2 controls were set up in one OTC-control plot pair in each of the three fences, making up an additional 12 collars.

### Data analysis

We tested the efficacy of the partitioning treatments in allowing/preventing the growth of roots and AMF into the collars, and their effect on soil microbial biomass carbon, using linear models with root dry weight, AMF ERM length or MBC as the response variable, and partitioning treatment as the predictor. The statistical similarity of soil respiration from the method control collars (C1 and C2) and the ‘soil without roots and AMF’ treatment was assessed using a linear mixed effects model (LMM). Partitioning treatment including the method controls and warming treatment were the fixed effects and collar nested within plots nested within fences were used as the random factor to account for repeated measures from the same collars. The R package *lme4* was used to build the mixed effects model, and *lmerTest* and *car* packages were used to assess the statistical significance of the fixed effects (Bates 2010; Bates and others 2014, Kuznetsova and others 2015; Bates and others 2017). All the analyses were conducted using R version 3.2.4 (The R Foundation for Statistical Computing, 2016).

### Results

The soil partitioning treatments were effective in allowing/preventing the growth of roots and AMF extraradical mycelium into the collars. Root biomass differed significantly between the partitioning treatments (*F* = 10.113, *df* = 2, *P* = 0.012), with ~10 times higher biomass (*P* = 0.008) in the ‘full soil’ treatment, at 1.22 ± 0.35 g, than in the other two treatments designed to keep out fine roots, which had 0.12 ± 0.01 g and 0.11 ± 0.03 g roots, respectively (Fig. S1a). The treatments also differed in AMF ERM levels, though treatment effect was not statistically significant (*F* = 3.8173, *df* = 2, *P* = 0.085). ‘Soil without roots and AMF’ treatment had the lowest ERM levels of 1.08 ± 0.95 m g^−1^ dry soil, differing significantly from the ‘soil without roots’ treatment (*P* = 0.033), which had the highest ERM levels at 3.56 ± 0.17 m g^−1^ dry soil, followed by the ‘full soil’ treatment with ERM of 2.23 ± 0.52 m g^−1^ dry soil (Fig. S1b). Partitioning treatment was not a statistically significant predictor for soil MBC (*F* = 4.3571, *df* = 2, *P* = 0.068). However, the ‘full soil’ treatment had the lowest MBC levels at 212.94 ± 25.05 mg kg^−1^ dry soil. This was significantly lower than the ‘soil without roots’ treatment (*P* = 0.026), which had the highest MBC levels at 325.68 ± 33.14 mg kg^−1^ dry soil, followed by the ‘soil without roots and AMF’ treatment that had 263.05 ± 21.70 mg kg^−1^ dry soil MBC (Fig. S1c).

Further, soil disturbance during collar setup did not influence soil respiration measurements. An LMM with all respiration collar treatments (the three partitioning treatments and the two method controls, C1 and C2) as the predictor, suggested that C1 and C2 were statistically indistinguishable from the ‘soil without roots and AMF’ treatment (*P* = 0.39 and 0.65, respectively).

**Fig. A.1.**
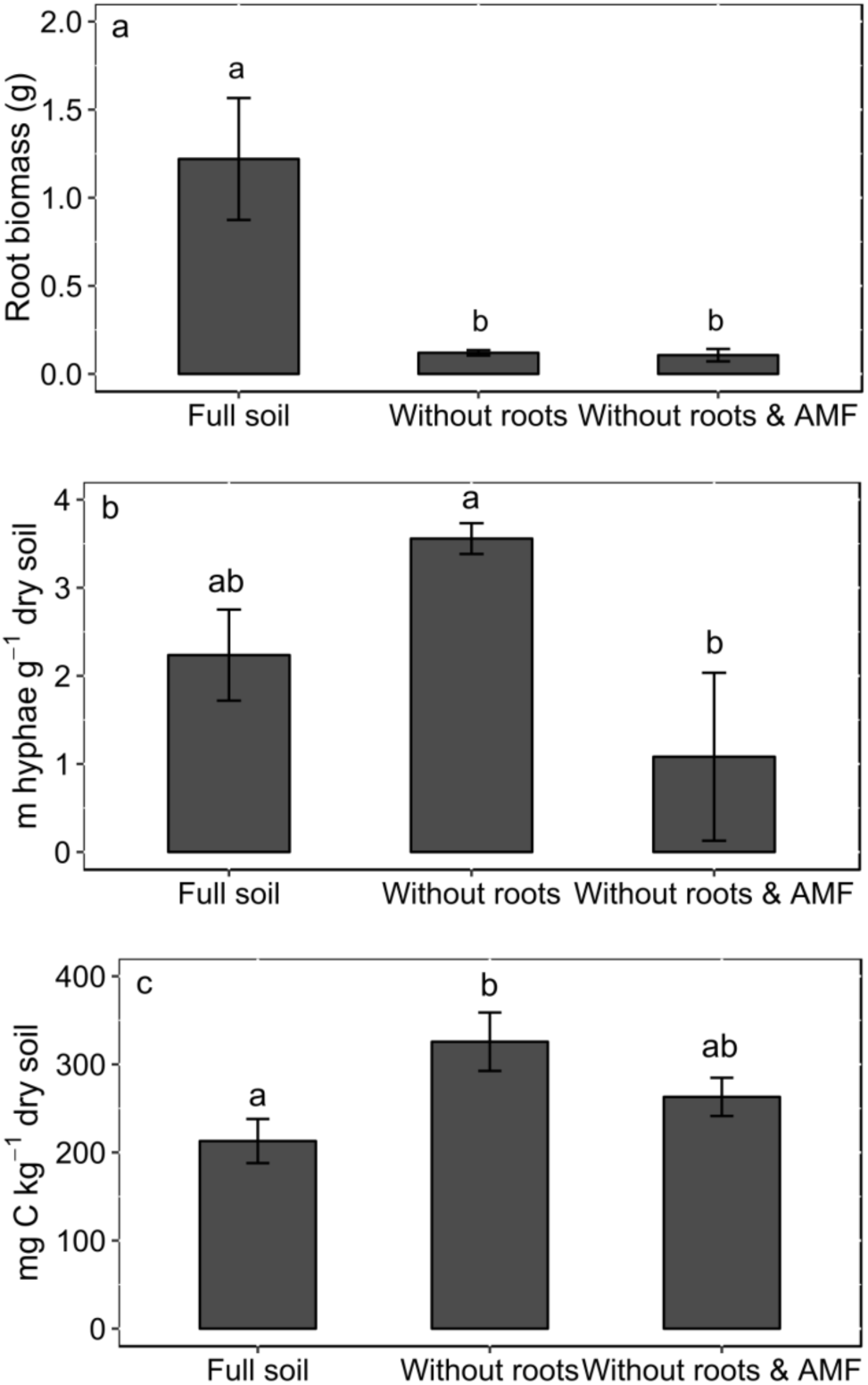
Estimates within each partitioning treatment for (a) root dry weight, (b) AMF extraradical mycelium length, and (c) microbial biomass carbon. Error bars are 1SE about the mean, with all values obtained from fixed effects statistics in the mixed effects models used for analyses. Different letters indicate statistically significant differences among treatments (*P* < 0.05).

## Supplemetary Information B: Warming effect on soil respiration per partitioning treatment

**Fig. B.1.**
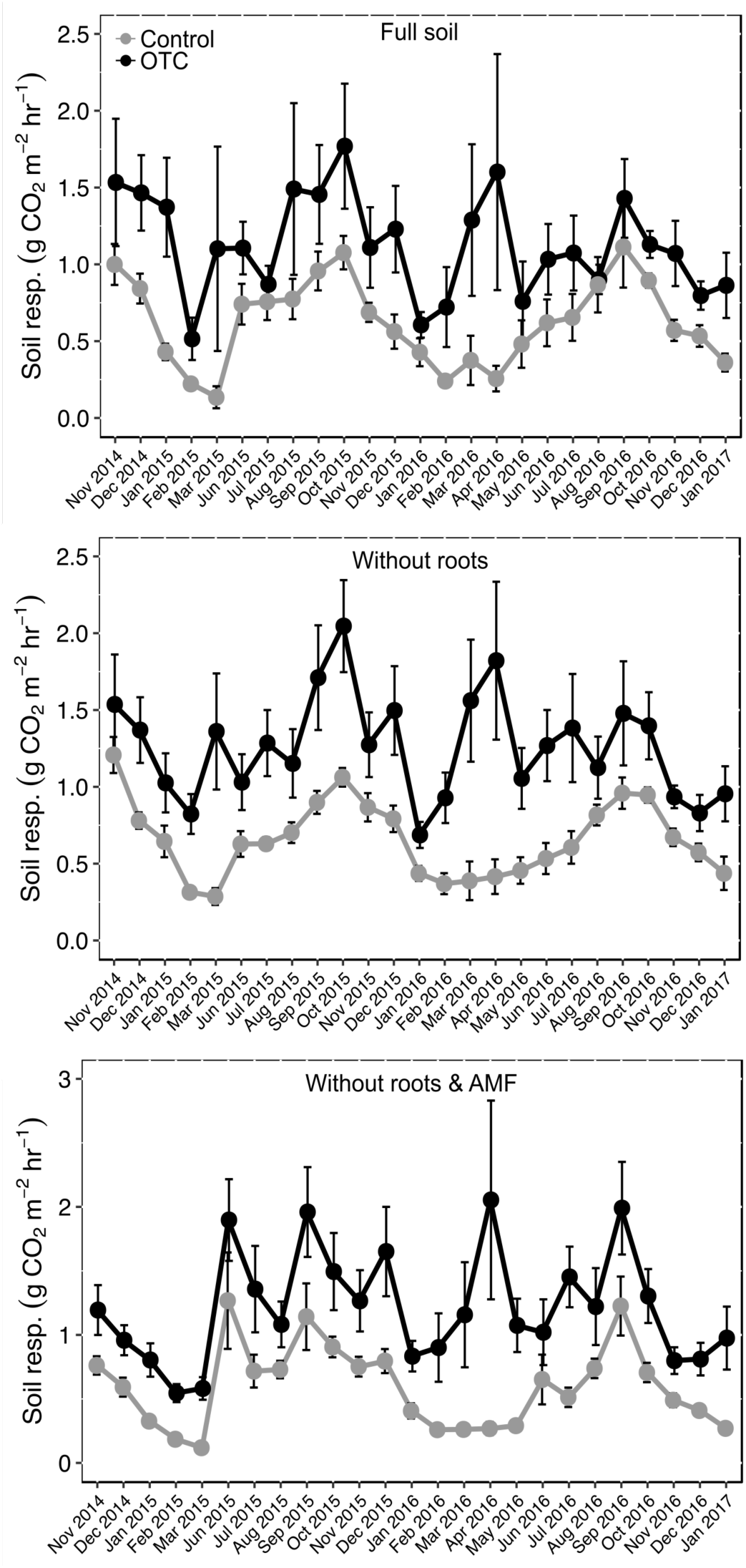
Average soil respiration in control and OTC plots averaged per month, in each of the partitioning treatments. Dots in grey represent control plots and those in black represent OTCs. Error bars represent 1SE around the mean.

## References

Arasumani M, Khan D, Das A, Lockwood I, Stewart R, Kiran RA, Muthukumar M, Bunyan M, Robin VV. 2018. Not seeing the grass for the trees: timber plantations and agriculture shrink tropical montane grassland by two-thirds over four decades in the Palani Hills, a Western Ghats Sky Island. PloS One 13:e0190003.

Aronson EL, McNulty SG. 2009. Appropriate experimental ecosystem warming methods by ecosystem, objective, and practicality. Agricultural and Forest Meteorology 149:1791–9.

Atkin OK, Edwards EJ, Loveys BR. 2000. Response of root respiration to changes in temperature and its relevance to global warming. New Phytologist 147:141–54.

Auffret MD, Karhu K, Khachane A, Dungait JAJ, Fraser F, Hopkins DW, Wookey PA, Singh BK, Freitag TE, Hartley IP, Prosser JI. 2016. The role of microbial community composition in controlling soil respiration responses to temperature. PloS One 11:e0165448.

Baayen RH, Davidson DJ, Bates DM. 2008. Mixed-effects modeling with crossed random effects for subjects and items. Journal of Memory and Language 59:390–412.

Baggs EM. 2006. Partitioning the components of soil respiration: a research challenge. Plant and Soil 284:1–5.

Bates D. 2010. lme4: Mixed-effects modeling with R. New York: Springer New York.

Bates D, Maechler M, Bolker B, Walker S, Christensen RHB, Singmann H, Dai B, Grothendieck G. 2014. Package ‘lme4’. Vienna: R Foundation for Statistical Computing.

Bates D, Maechler M, Bolker B, Walker S, Christensen RHB, Singmann H, Dai B, Grothendieck G, Green P. 2017. Package ‘lme4’. Vienna: R Foundation for Statistical Computing.

Birgander J, Rousk J, Olsson PA. 2017. Warmer winters increase the rhizosphere carbon flow to mycorrhizal fungi more than to other microorganisms in a temperate grassland. Global Change Biology 23:5372–82.

Bond-Lamberty B, Wang C, Gower ST. 2004. A global relationship between the heterotrophic and autotrophic components of soil respiration? Global Change Biology 10:1756–66.

Bond-Lamberty B, Thomson A. 2010. Temperature-associated increases in the global soil respiration record. Nature 464:579–82.

Bronson DR, Gower ST, Tanner M, Linder S, Van Herk I. 2007. Response of soil surface CO_2_ flux in a boreal forest to ecosystem warming. Global Change Biology 14:856–67.

Buchmann N. 2000. Biotic and abiotic factors controlling soil respiration rates in Picea abies stands. Soil Biology and Biochemistry 32:1625–35.

Cartmill AD. 2011. Effect of warming and precipitation distribution on soil respiration and mycorrhizal abundance in post oak savannah. Texas A&M University.

Classen AT, Sundqvist MK, Henning JA, Newman GS, Moore JAM, Cregger MA, Moorhead LC, Patterson CM. 2015. Direct and indirect effects of climate change on soil microbial and soil microbial-plant interactions: What lies ahead? Ecosphere 6:art130.

Conant RT, Dalla-Betta P, Klopatek CC, Klopatek JM. 2004. Controls on soil respiration in semiarid soils. Soil Biology and Biochemistry 36:945–51.

Cunnings I, Finlayson I. 2015. Mixed effects modeling and longitudinal data analysis. In: Plonsky L, editor. Advancing quantitative methods in second language research. Second Language Acquisition Research Series. New York: Routledge, Taylor and Francis Group. p159–81.

DeAngelis KM, Pold G, Topçuoğlu BD, van Diepen LTA, Varney RM, Blanchard JL, Melillo J, Frey SD. 2015. Long-term forest soil warming alters microbial communities in temperate forest soils. Frontiers in Microbiology 6.

Godfree R, Robertson B, Bolger T, Carnegie M, Young A. 2011. An improved hexagon open-top chamber system for stable diurnal and nocturnal warming and atmospheric carbon dioxide enrichment. Global Change Biology 17:439–51.

Gunatilleke N, Pethiyagoda R, Gunatilleke S. 2008. Biodiversity of Sri Lanka. Journal of the National Science Foundation of Sri Lanka 36:25–61.

Hanson PJ, Edwards NT, Garten CT, Andrews JA. 2000. Separating root and soil microbial contributions to soil respiration: A review of methods and observations. Biogeochemistry 48:115–146.

Hashimoto S, Carvalhais N, Ito A, Migliavacca M, Nishina K, Reichstein M. 2015. Global spatiotemporal distribution of soil respiration modeled using a global database. Biogeosciences 12:4121–32.

Hawkes CV, Hartley IP, Ineson P, Fitter AH. 2008. Soil temperature affects carbon allocation within arbuscular mycorrhizal networks and carbon transport from plant to fungus. Global Change Biology 14:1181–90.

Heinemeyer A, Ineson P, Ostle N, Fitter AH. 2006. Respiration of the external mycelium in the arbuscular mycorrhizal symbiosis shows strong dependence on recent photosynthates and acclimation to temperature. New Phytologist 171:159–70.

Heinemeyer A, Wilkinson M, Vargas R, Subke J-A, Casella E, Morison JIL, Ineson P. 2012. Exploring the ‘overflow tap’ theory: linking forest soil CO_2_ fluxes and individual mycorrhizosphere components to photosynthesis. Biogeosciences 9:79–95.

Hoover DL, Knapp AK, Smith MD. 2016. The immediate and prolonged effects of climate extremes on soil respiration in a mesic grassland. Journal of Geophysical Research: Biogeosciences 121:1034–44.

Kelting DL, Burger JA, Edwards GS. 1998. Estimating root respiration, microbial respiration in the rhizosphere, and root-free soil respiration in forest soils. Soil Biology and Biochemistry 30:961–968.

Kim Y-C, Gao C, Zheng Y, He X-H, Yang W, Chen L, Wan S-Q, Guo L-D. 2015. Arbuscular mycorrhizal fungal community response to warming and nitrogen addition in a semiarid steppe ecosystem. Mycorrhiza 25:267–76.

Kirschbaum MUF. 2004. Soil respiration under prolonged soil warming: are rate reductions caused by acclimation or substrate loss? Global Change Biology 10:1870–7.

Kotze DJ, Samways MJ. 2001. No general edge effects for invertebrates at Afromontane forest/grassland ecotones. Biodiversity and Conservation 10:443–466.

Kuznetsova A, Brockhoff PB, Christensen RHB. 2015. Package ‘lmerTest’. Vienna: R Foundation for Statistical Computing.

Lefcheck J. 2016. Package ‘piecewiseSEM’. Vienna: R Foundation for Statistical Computing.

Li D, Zhou X, Wu L, Zhou J, Luo Y. 2013. Contrasting responses of heterotrophic and autotrophic respiration to experimental warming in a winter annual-dominated prairie. Global Change Biology 19:3553–64.

Li Y, Zhou G, Huang W, Liu J, Fang X. 2016. Potential effects of warming on soil respiration and carbon sequestration in a subtropical forest. Plant and Soil 409:247–57.

Liu T, Xu Z-Z, Hou Y-H, Zhou G-S. 2016. Effects of warming and changing precipitation rates on soil respiration over two years in a desert steppe of northern China. Plant and Soil 400:15–27.

Lu M, Zhou X, Yang Q, Li H, Luo Y, Fang C, Chen J, Yang X, Li B. 2013. Responses of ecosystem carbon cycle to experimental warming: a meta-analysis. Ecology 94:726–38.

Luo Y, Wan S, Hui D, Wallace LL. 2001. Acclimatization of soil respiration to warming in a tall grass prairie. Nature 413:622–5.

Luo C, Rodriguez-R LM, Johnston ER, Wu L, Cheng L, Xue K, Tu Q, Deng Y, He Z, Shi JZ, Yuan MM, Sherry RA, Li D, Luo Y, Schuur EAG, Chain P, Tiedje JM, Zhou J, Konstantinidis KT. 2014. Soil microbial community responses to a decade of warming as revealed by comparative metagenomics. Applied and Environmental Microbiology 80:1777–86.

Marthews TR, Riutta T, Oliveras Menor I, Urrutia R, Moore S, Metcalfe D, Malhi Y, Phillips O, Huaraca Huasco W, Ruiz Jaén M, Girardin C, Butt N, Cain R, colleagues from the Rainfor and GEM networks. 2014. Measuring tropical forest carbon allocation and cycling: A RAINFOR-GEM field manual for intensive census plots (v3.0).

Mayer M, Sandén H, Rewald B, Godbold DL, Katzensteiner K. 2017. Increase in heterotrophic soil respiration by temperature drives decline in soil organic carbon stocks after forest windthrow in a mountainous ecosystem. Functional Ecology 31:1163–72.

Melillo JM, Steudler PA, Aber JD, Newkirk K, Lux H, Bowles FP, Catricala C, Magill A, Ahrens T, Morrisseau S. 2002. Soil warming and carbon-cycle feedbacks to the climate system. Science 298:2173–6.

Metcalfe DB, Fisher RA, Wardle DA. 2011. Plant communities as drivers of soil respiration: pathways, mechanisms, and significance for global change. Biogeosciences 8:2047–61.

Moyano FE, Manzoni S, Chenu C. 2013. Responses of soil heterotrophic respiration to moisture availability: An exploration of processes and models. Soil Biology and Biochemistry 59:72–85.

Nakagawa S, Schielzeth H. 2013. A general and simple method for obtaining R^2^ from generalized linear mixed-effects models. Methods in Ecology and Evolution 4:133–42.

Olsson P, Linder S, Giesler R, Hogberg P. 2005. Fertilization of boreal forest reduces both autotrophic and heterotrophic soil respiration. Global Change Biology 11:1745–53.

Orchard VA, Cook FJ. 1983. Relationship between soil respiration and soil moisture. Soil Biology and Biochemistry 15:447–53.

Overbeck G, Muller S, Fidelis A, Pfadenhauer J, Pillar V, Blanco C, Boldrini I, Both R, Forneck E. 2007. Brazil’s neglected biome: The South Brazilian Campos. Perspectives in Plant Ecology, Evolution and Systematics 9:101–16.

Parr CL, Lehmann CER, Bond WJ, Hoffmann WA, Andersen AN. 2014. Tropical grassy biomes: misunderstood, neglected, and under threat. Trends in Ecology and Evolution 29:205–13.

Raich JW, Schlesinger WH. 1992. The global carbon dioxide flux in soil respiration and its relationship to vegetation and climate. Tellus 44:81–99.

Robin VV, Nandini R. 2012. Shola habitats on sky islands: status of research on montane forests and grasslands in southern India. Current Science. 103:1427–37.

Rodeghiero M, Cescatti A. 2006. Indirect partitioning of soil respiration in a series of evergreen forest ecosystems. Plant and Soil 284:7–22.

Romero-Olivares AL, Allison SD, Treseder KK. 2017. Soil microbes and their response to experimental warming over time: A meta-analysis of field studies. Soil Biology and Biochemistry 107:32–40.

Rudgers JA, Kivlin SN, Whitney KD, Price MV, Waser NM, Harte J. 2014. Responses of highaltitude graminoids and soil fungi to 20 years of experimental warming. Ecology 95:1918–1928.

Rustad L, Campbell JL, Marion G, Norby R, Mitchell M, Hartley A, Cornelissen J, Gurevitch J, Global Change and Terrestrial Ecosystems Network of Ecosystem Warming Studies. 2001. A meta-analysis of the response of soil respiration, net nitrogen mineralization, and aboveground plant growth to experimental ecosystem warming. Oecologia 126:543–62.

Savage K, Davidson EA, Richardson AD. 2008. A conceptual and practical approach to data quality and analysis procedures for high-frequency soil respiration measurements. Functional Ecology 22:1000–7.

Schindlbacher A, Zechmeister-Boltenstern S, Jandl R. 2009. Carbon losses due to soil warming: Do autotrophic and heterotrophic soil respiration respond equally? Global Change Biology 15:901–13.

Schindlbacher A, Rodler A, Kuffner M, Kitzler B, Sessitsch A, Zechmeister-Boltenstern S. 2011. Experimental warming effects on the microbial community of a temperate mountain forest soil. Soil Biology and Biochemistry 43:1417–25.

Scott-Denton LE, Rosenstiel TN, Monson RK. 2006. Differential controls by climate and substrate over the heterotrophic and rhizospheric components of soil respiration. Global Change Biology 12:205–16.

Singh BK, Bardgett RD, Smith P, Reay DS. 2010. Microorganisms and climate change: terrestrial feedbacks and mitigation options. Nature Reviews Microbiology 8:779–90.

Sukumar R, Suresh HS, Ramesh R. 1995. Climate change and its impact on tropical montane ecosystems in southern India. Journal of Biogeography 22:533–6.

Treseder KK, Marusenko Y, Romero-Olivares AL, Maltz MR. 2016. Experimental warming alters potential function of the fungal community in boreal forest. Global Change Biology 22:3395–404.

Wang X, Liu L, Piao S, Janssens IA, Tang J, Liu W, Chi Y, Wang J, Xu S. 2014. Soil respiration under climate warming: differential response of heterotrophic and autotrophic respiration. Global Change Biology 20:3229–37.

Wangdi N, Mayer M, Nirola MP, Zangmo N, Orong K, Ahmed IU, Darabant A, Jandl R, Gratzer G, Schindlbacher A. 2017. Soil CO_2_ efflux from two mountain forests in the eastern Himalayas, Bhutan: components and controls. Biogeosciences 14:99–110.

Wardle DA, Bardgett RD, Klironomos JN, Setälä H, van der Putten WH, Wall D. 2004. Ecological linkages between aboveground and belowground biota. Science 304:1629–33.

Wei W, Weile C, Shaopeng W. 2010. Forest soil respiration and its heterotrophic and autotrophic components: Global patterns and responses to temperature and precipitation. Soil Biology and Biochemistry 42:1236–1244.

Wu Z, Dijkstra P, Koch GW, Peñuelas J, Hungate BA. 2011. Responses of terrestrial ecosystems to temperature and precipitation change: a meta-analysis of experimental manipulation. Global Change Biology 17:927–42.

Yan L, Chen S, Huang J, Lin G. 2009. Differential responses of auto- and heterotrophic soil respiration to water and nitrogen addition in a semiarid temperate steppe. Global Change Biology 16:2345–57.

Zogg GP, Zak DR, Ringelberg DB, White DC, MacDonald NW, Pregitzer KS. 1997. Compositional and functional shifts in microbial communities due to soil warming. Soil Science Society of America Journal 61:475.

Zuur AF, editor. 2009. Mixed effects models and extensions in ecology with R. New York: Springer.

## References

Anderson JPE, Domsch KH. 1978. A physiological method for the quantitative measurement of microbial biomass in soils. Soil Biology & Biochemistry 10:215–221.

Bates DM. 2010. lme4: Mixed-effects modeling with R. New York: Springer New York.

Brundrett M, Melville L, Peterson L, editors. 1994. Practical methods in mycorrhiza research. Mycologue Publications.

Fanin N, Hättenschwiler S, Barantal S, Schimann H, Fromin N. 2011. Does variability in litter quality determine soil microbial respiration in an Amazonian rainforest? Soil Biology & Biochemistry 43:1014–22.

Höper H. 2006. Substrate induced respiration. In: Bloem J, Hopkins DW, Benedetti A, editors. Microbiological methods for assessing soil quality. Wallingford, UK?; Cambridge, MA: CABI Pub. p 84–92.

